# Canonical mTOR signaling supports complete fin regeneration

**DOI:** 10.64898/2026.03.27.714790

**Authors:** Josane F. de Sousa, Gabriela Lima, Louise Perez, Michaela Tsanova, Cyrus Bronson, Garrison Boehl, Icyss Sargeant, Rogerio Gomes, Aline C. Dragalzew, Wainna B. Mendes, Igor Schneider

## Abstract

The mechanistic target of rapamycin (mTOR) pathway is a deeply conserved regulator of cellular growth, metabolism, and tissue repair. In vertebrates, mTOR activity is broadly required for wound healing and regeneration. Recent work in salamanders identified unique amino acid expansions within the mTOR protein. These expansions are thought to confer hypersensitive kinase activity, which was proposed to support rapid translational activation and exceptional limb regeneration capacity. This has raised the possibility that canonical, mammalian-like mTOR signaling may be insufficient to support regeneration of complex appendages. Here, we test this hypothesis by investigating fin regeneration in the Senegal bichir (*Polypterus senegalus*), a ray-finned fish capable of fully regenerating fins containing bone, cartilage, muscle, and connective tissue with cellular complexity comparable to tetrapod limbs. We show that fin amputation triggers rapid activation of canonical mTOR signaling, and pharmacological inhibition of mTOR with rapamycin abolishes fin regeneration despite successful wound closure. Single-nucleus RNA sequencing (snRNA-seq) across multiple regenerative stages reveals that mTOR pathway components are selectively upregulated in proliferative epidermal and connective tissue cells, and myeloid cell populations. Rather than causing a global shutdown of gene expression, mTOR inhibition specifically dampened the induction of translational machinery and glycolytic programs, two canonical downstream outputs of mTOR signaling. In addition, myeloid cells exhibited pronounced sensitivity to mTOR inhibition, showing attenuated immune-competent and pro-regenerative programs such as interferon-associated signaling, antigen processing, and innate immune coordination. We propose that salamander-specific mTOR hypersensitivity represents an evolutionary acceleration of an ancestral regenerative program, rather than a prerequisite for limb regeneration.

**Significance Statement:** The ability to regenerate complex appendages varies widely among vertebrates, yet the molecular basis of this variation remains poorly understood. Salamanders harbor a uniquely hypersensitive form of mTOR that has been proposed to be essential for limb regeneration. Using *Polypterus* fin regeneration as a model system, we demonstrate that a canonical vertebrate mTOR protein is compatible with regeneration of a complex, limb-like appendage. Our results show that mTOR acts through selective metabolic, translational, and immune programs. These findings suggest that salamander-specific mTOR hypersensitivity represents a derived enhancement rather than a condition for appendage regeneration and provide insight into the evolutionary foundations of regenerative competence.

## Introduction

The mTOR signaling pathway is a deeply conserved eukaryotic network that integrates intracellular and environmental cues to regulate protein and nucleotide synthesis, cell growth and proliferation, survival, and migration (1-3). The central component of this signaling pathway is mTOR, a serine/threonine kinase originally identified as the direct target of rapamycin, a potent antifungal compound with immunosuppressive and antitumor activities. Through its assembly into functionally distinct complexes, mTOR coordinates cellular anabolic and metabolic responses with organismal physiology.

Across vertebrates, mTOR signaling has been broadly implicated in tissue repair and regeneration in a wide range of organs and contexts. In mammals, mTOR activity is required for regeneration and repair of various tissue and cell types, such as skeletal muscle (4, 5), neurons (6), hair follicles (7), and skin (8). In non-mammalian vertebrates, mTOR is necessary for regeneration of complex structures, including tail regeneration in geckos (9), tail fin (10-12), retinal pigment epithelium (13), and heart regeneration (14) in zebrafish, tail (15) and spinal cord (16) regeneration in *Xenopus laevis*, and limb regeneration in axolotls (17, 18). Together, these studies establish mTOR as a central and evolutionarily conserved regulator of regenerative processes across vertebrates.

Recent work has shown that salamanders harbor unique amino acid expansions within the mTOR protein that render the kinase primed for activation (17). In addition, axolotls have been shown to mount a systemic response to injury involving widespread cell-cycle re-entry in an mTOR-dependent manner (18). Together, these findings suggest that salamanders have evolved a response to injury mediated in part by an enhanced mTOR protein. Whether such salamander-specific mTOR hypersensitivity is required for regeneration of complex appendages, or whether canonical vertebrate mTOR signaling is sufficient, remains unresolved.

To address this, we investigated the role of mTOR signaling during pectoral fin regeneration of the early-diverging bony fish *Polypterus senegalus* (Fig. 1*A*). This system offers important advantages relative to zebrafish fin ray regeneration. In contrast to caudal fin rays, pectoral fins are the direct homologues of limbs in fishes (19). *Polypterus* also deploys a conserved cellular and transcriptional program during fin regeneration that parallels key aspects of salamander limb regeneration (20, 21). Moreover, *Polypterus* regrows the entire fin (22), including the most proximal, endoskeleton-bearing region, which shares with limbs the complex cellular composition (muscle, tendons, joints, chondrocytes, nerves etc.), not found in fin rays (23). Finally, *Polypterus* diverged from teleosts prior to the teleost-specific whole genome duplication (Fig. 1*B*), and as a result, share with salamanders a 1:1 gene orthology, which facilitates comparisons of orthologous genes (24).

**Fig. 1.**
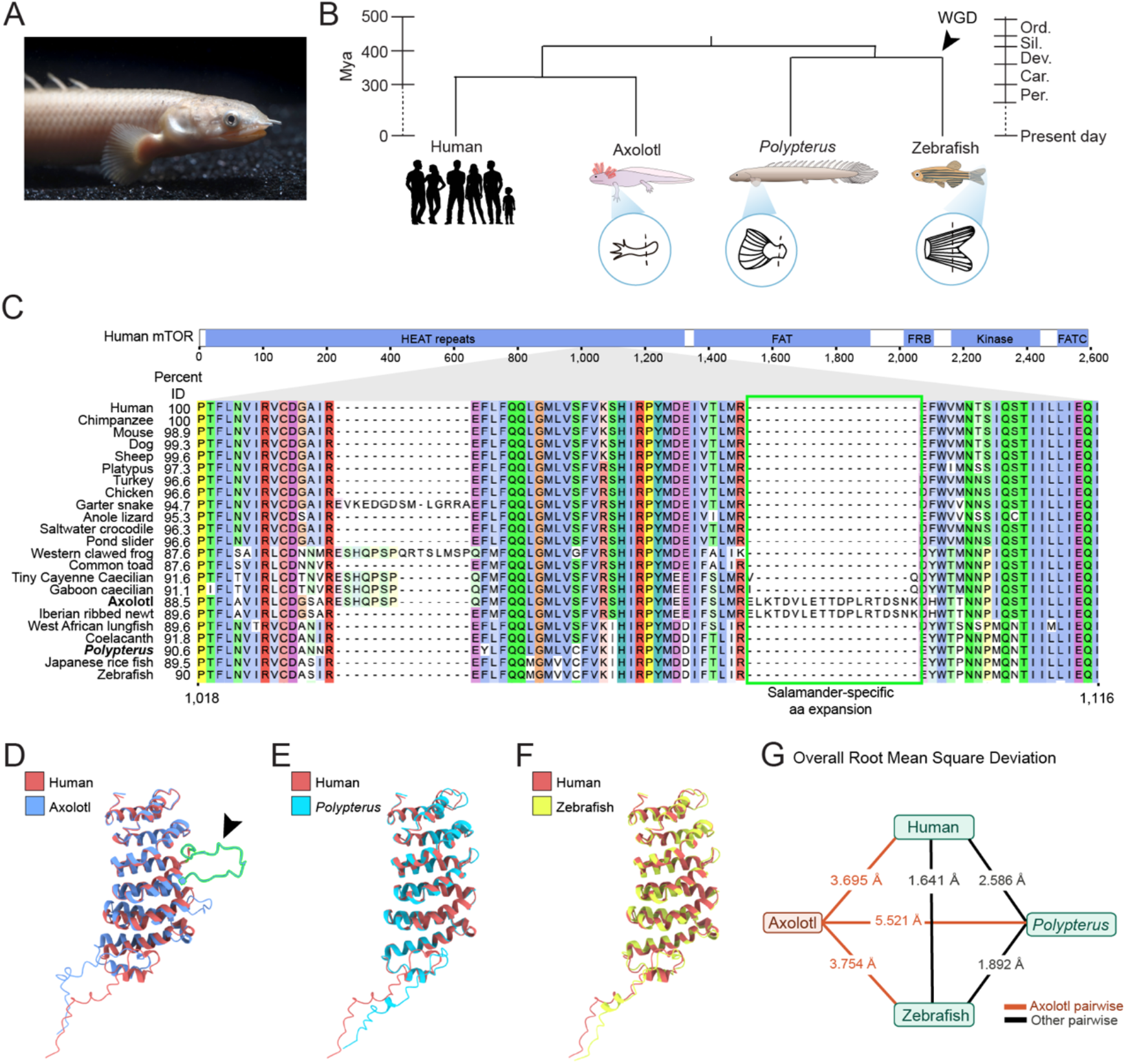
*Polypterus* harbors a canonical mTOR protein. (A) Photo of a Polypterus individual. (B) Phylogenetic relationships between human, axolotl, Polypterus and zebrafish (C) Schematic representation of the human mTOR protein, its domains, and the region within the HEAT repeat harboring the salamander-specific amino acid expansion (green box); percent identities of full-length mTOR proteins relative to the human mTOR are shown for 22 vertebrates. (D-F) Pairwise structural alignment of 250 residues of the human mTOR protein spanning the HEAT repeat region harboring the salamander-specific expansion are shown for human and axolotl (D), human and *Polypterus* (E), and human and zebrafish (F). (G) Overall Root Mean Square Deviation obtained from pairwise structural comparisons between axolotl, human, *Polypterus* and zebrafish mTOR HEAT repeat region. WGD, whole genome duplication; aa, amino acid; Mya; million years ago.

Here, we demonstrate that the *Polypterus* mTOR, which lacks the amino acid expansions found in axolotls, is required for fin regeneration. mTOR signaling activation occurred within hours after fin amputation and is sustained throughout regeneration. Pharmacological inhibition of mTOR signaling with rapamycin or INK128 does not prevent wound closure but blocks regenerative outgrowth. Analysis of snRNA-seq across fin regeneration stages identifies cycling epidermal and connective tissue cells, as well as myeloid cells, as primary sites of expression of mTOR pathway gene expression. Inhibition of mTOR signaling suppresses the expression of genes involved in protein translation and glycolytic metabolism, two canonical downstream programs regulated by mTOR activity. Finally, mTOR inhibition disrupts interferon and antigen presentation programs in myeloid cells, consistent with a model in which mTOR activity preferentially supports immune programs linked to regenerative coordination and tissue resolution. Collectively, our findings suggest that mTOR dependence in appendage regeneration likely reflects an ancestral vertebrate state and that salamander-specific mTOR hypersensitivity represents a derived acceleration of this program rather than the origin of limb regenerative competence.

## Results

### *Polypterus* harbors a canonical mTOR protein

To determine whether *Polypterus* mTOR exhibits lineage-specific features, we performed multispecies alignments of mTOR protein sequences from *Polypterus* and 23 additional vertebrates spanning cartilaginous fishes to mammals. The full alignment is publicly available via Figshare (DOI: 10.6084/m9.figshare.31120030). Across vertebrates, mTOR is highly conserved, with overall sequence identity ranging from 87.6% to 98.9%, and *Polypterus* does not display conspicuous insertions or deletions relative to other non-salamander taxa (Fig. 1C). In contrast, the previously described salamander-specific amino acid expansion, implicated in increased mTOR sensitivity, is localized within the HEAT repeat region (Fig. 1C). Consistent with this, pairwise comparisons of axolotl, *Polypterus*, and zebrafish mTOR against human mTOR revealed that the HEAT repeat region (human mTOR residues 1-1380) locates to a variable portion of the protein (86.7-94.8% identity), whereas key functional domains are markedly conserved (FRB: 97-99%; kinase domain: 98-99%) (Dataset S1). Focusing on the region encompassing the salamander-specific expansion (human mTOR residues 990-1150), sequence identity dropped further (86-95%), highlighting this segment as a potential hotspot of divergence within an otherwise highly conserved protein (Dataset S1).

To assess whether these sequence differences translate into structural divergence, we generated three-dimensional models of this HEAT repeat region for human, axolotl, *Polypterus*, and zebrafish using ColabFold (25), an optimized implementation of AlphaFold2 (26). Structural alignment to the human model revealed that the axolotl-specific expansion forms an unstructured loop that protrudes from the conserved core (Fig. 1D), whereas *Polypterus* and zebrafish closely overlap with the human structure (Fig. 1E,F). Quantification using root mean square deviation (RMSD) supported this observation: axolotl comparisons exhibited substantially higher RMSD values (3.7-5.5 Å), while pairwise comparisons among human, *Polypterus*, and zebrafish remained low (1.6-2.6 Å) (Fig. 1G). Together, these analyses demonstrate that *Polypterus* mTOR retains the conserved vertebrate architecture and lacks the salamander-specific insertion associated with enhanced kinase sensitivity. Both sequence and structural evidence support that *Polypterus* mTOR is highly similar to mammalian and teleost mTOR. We therefore classify *Polypterus* mTOR as canonical.

### *Polypterus* mTOR pathway is rapidly activated upon injury and required for fin regeneration

To determine whether the canonical *Polypterus* mTOR is capable to rapidly activate the mTOR signaling, we assessed mTOR activity during *Polypterus* fin regeneration via phosphorylation of ribosomal protein S6 (pS6) immunofluorescence. As soon as 3 hours post-amputation (hpa), we detected pS6 signal in cells contributing to epithelial cells covering the wound (Fig. 2*A*). From 18 hpa to 3 days post-amputation (dpa), as the wound epidermis thickened, pS6 signal increased and was also visible in subjacent mesenchymal cells. This pattern was maintained across 5 and 9 dpa stages. To confirm our observations of pS6 flourescence signal, we performed assessed pS6^Ser240/244^ to S6 protein ratios normalized to β-actin across regeneration stages. Concordantly, we observed increased pS6/S6 ratios as soon as 2 hpa, which remained sustained at 1 and 3 dpa (Fig 2.*B*, Dataset S2). Therefore, as seen in axolotls, mTOR signaling is rapidly activated upon *Polypterus* fin amputation.

**Fig. 2.**
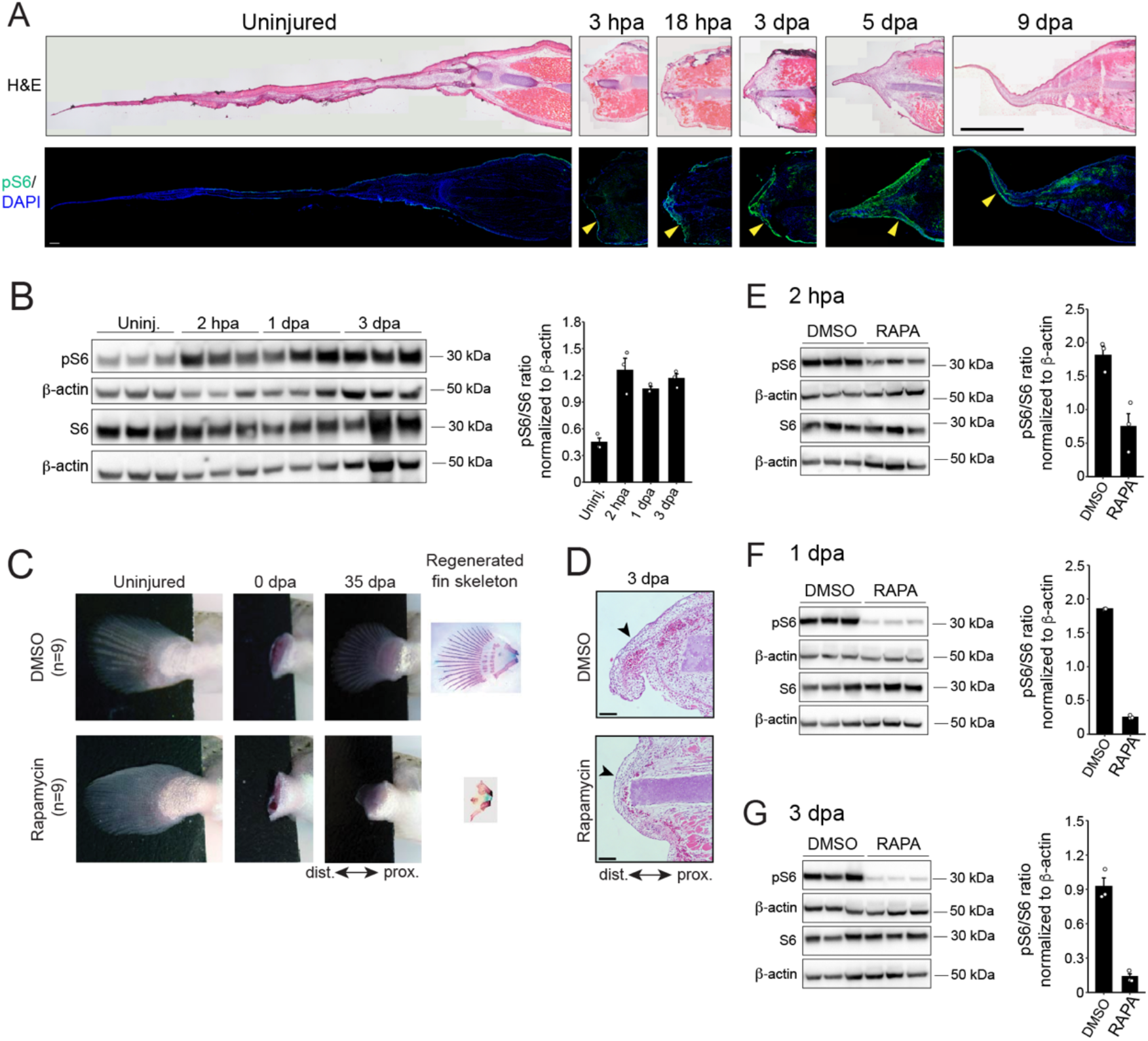
mTOR signaling is rapidly activated and required for *Polypterus* fin regeneration. (A) Top: Hematoxylin and Eosin (H&E) staining; Bottom: pS6 (green) immunofluorescence and nuclei counterstaining with DAPI (blue) in uninjured and regenerating fins; yellow arrowheads denote wound epidermis. (B) Left: Western blots for pS6, S6 and β-actin in uninjured and regenerating fins; Right: Quantification of pS6/S6 ratios normalized relative to β-actin. (C) Uninjured, 0 dpa and 35 dpa fins and regenerated fin skeleton of DMSO-(top) and rapamycin-(bottom) treated animals. (D) H&E-stained fin sections of the wound epidermis of DMSO- and rapamycin-treated animals at 3 dpa; black arrowheads denote wound epidermis. (E-G) Left: Western blots for pS6, S6 and β-actin in DMSO- and rapamycin-treated animals at 2 hpa (E), 1 dpa (F) and (G); Right: corresponding quantification of pS6/S6 ratios normalized relative to β-actin. Scale bars: 100 μM.

To determine if mTOR signaling is required for fin regeneration, we treated *Polypterus* with rapamycin, a potent inhibitor of the mTOR signaling pathway (27). Treatment consisted of immersing animals immediately after fin amputation in system water containing either dimethyl sulfoxide (DMSO) or rapamycin (2.5 μM diluted in DMSO), with treatment solutions replaced daily until regeneration was assessed. In contrast to DMSO-treated controls, rapamycin-treated animals consistently failed to regrow fins after amputation (Fig. 2*C*, *SI Appendix*, Fig S1*A*). To independently validate these findings, we treated animals with INK128, a next generation mTOR inhibitor recently shown to impair axolotl limb regeneration (17). Whereas rapamycin acts as an allosteric inhibitor, INK128 is an ATP-competitive inhibitor that blocks the catalytic activity of mTOR. Treatment with INK128 (0.5 μM) severely impaired fin regeneration (*SI Appendix*, Fig S1*B*). Histological analysis of DMSO- and rapamycin-treated fins revealed that mTOR inhibition did not prevent early wound closure, despite preventing fin outgrowth (Fig. 2*D*). To confirm effective inhibition of mTOR signaling by rapamycin, we assessed pS6/S6 ratios via western blotting at 2 hpa, 1 dpa, and 3 dpa. Our experiments revealed a significant decrease in pS6/S6 ratios at 2 hpa upon rapamycin treatment (Fig. 2*E*), which was accentuated at 1 dpa (Fig. 2*F*) and 3 dpa (Fig. 2*G*, Dataset S2). Together, these results demonstrate that mTOR signaling is rapidly activated following fin amputation and is specifically required for subsequent regenerative outgrowth.

### Cell-type-specific upstream regulation of mTOR activity during fin regeneration

Previous work established that mTOR activity is required for successful axolotl limb regeneration, where it promotes a rapid translational upregulation of mRNAs, especially those encoding components of the protein synthesis machinery (17). These translational responses were most prominent in intermediate and basal wound epidermis, as well as in immune cell populations. Nevertheless, the downstream consequences of mTOR inhibition at cellular resolution remain incompletely characterized, including which regenerative cell populations are most affected, and which key signaling pathways and biological processes are altered in response to mTOR inhibition.

To gain mechanistic insight into how mTOR signaling regulates fin regeneration, we examined mTOR pathway activity in *Polypterus* by identifying cells expressing mTOR components, downstream effectors, and candidate upstream regulators. To this end, we generated a snRNA-seq dataset from rapamycin-treated *Polypterus* fins collected at regenerative stages matched to a previously published snRNA-seq dataset of DMSO-treated fins (21).

Together, these datasets comprise approximately 116,000 nuclei from uninjured fins and from DMSO- or rapamycin-treated fins at 1, 3, and 7 dpa, with biological replicates for each condition (*SI Appendix*, Fig. S2*A-E*). Unsupervised clustering identified 36 transcriptionally distinct clusters, whose cell type identities were inferred based on marker gene expression and prior annotation (21) (Fig. 3*A*, *SI Appendix*, Fig. S2*F*). All clusters were represented in both DMSO- and rapamycin-treated samples. To facilitate downstream analyses, we consolidated these clusters into 14 broader cell classes, including epidermal subtypes (proliferating, basal, intermediate, superficial, and specialized), erythrocytes, platelets, endothelial cells, lymphocytes, myeloid cells, glial cells, muscle, connective tissue (CT), and proliferative CT cells (Fig. 3*B*). Then we evaluated expression of mTOR pathway genes during the normal course of fin regeneration in DMSO-treated samples.

**Fig. 3.**
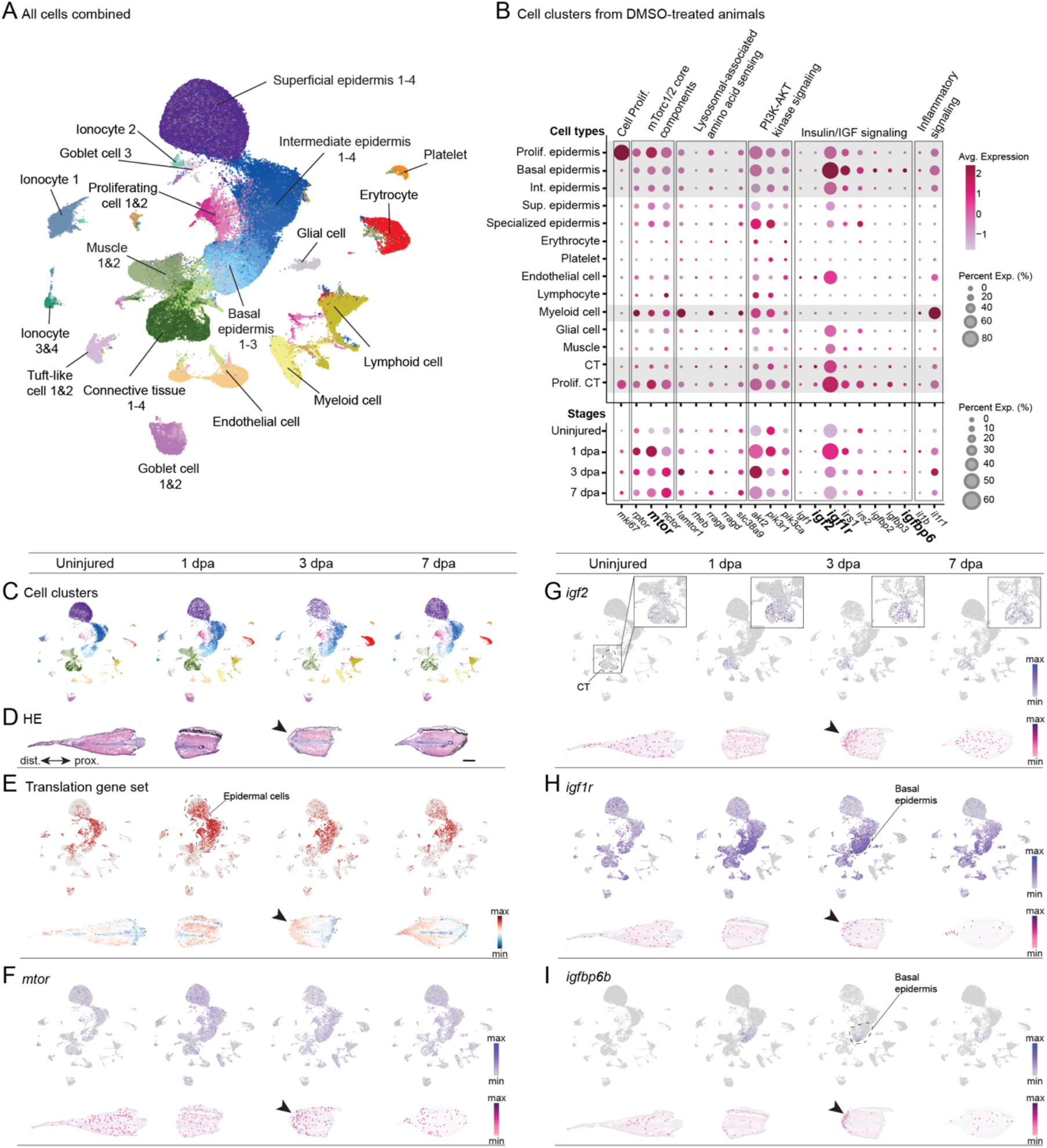
SnRNA-seq reveal upstream regulators and downstream targets of mTOR signaling during fin regeneration. (A) UMAP showing combined cell clusters from all stages of fin regeneration in DMSO- and rapamycin-treated animals. (B) Dot plot showing expression of mTOR signaling components and modulators in consolidated cell clusters and across fin regeneration stages in DMSO-treated animals. (C) UMAP of cell clusters across fin regeneration stages in DMSO-treated animals. (D) Histological sections of *Polypterus* fins during homeostasis and regeneration stages. (E) UMAP (top) and spatial transcriptomic (bottom) sections showing enrichment of a translational machinery set during homeostasis and regeneration stages. (F-I) UMAP (top) and spatial transcriptomic (bottom) sections showing expression of *mtor* (F), *igf2* (G), *igf1r* (H) and *igfbp6b* (I) during homeostasis and regeneration stages; insets (G) show zoomed in view of connective tissue (CT); basal epidermis is outlined (H and I); black arrowheads (D-I) denote wound epidermis. Scale bar = 1 cm (D).

We first examined expression of core mTOR pathway components (*mtor, rptor*, and *rictor*) to define where upstream regulatory signals might converge. Consistent with previous findings of mTOR activity in cycling cells during axolotl limb regeneration (18), transcripts encoding mTorc1 and mTorc2 core components were upregulated during regenerative stages (Fig. 3*B*), and was most pronounced in proliferative epidermal and CT cells (marked by *mki67* expression). As seen in axolotls (17), expression of mTorc1/2 core components was also evident across epidermal cell subtypes and myeloid cells. We next assessed expression of genes representing four major upstream regulatory axes known to converge on mTOR signaling: lysosome-associated amino acid sensing, PI3K-AKT kinase signaling, insulin/IGF signaling, and inflammatory signaling. The lysosome-associated amino acid sensing genes *lamtor1, rraga*, and *slc38a9*, were most highly expressed in myeloid cells. This pattern is consistent with increased lysosomal activity associated with debris clearance following injury, while also suggesting enhanced amino acid-dependent mTOR regulation in myeloid cells. Components of the PI3K-AKT pathway (*pik3ca, pik3r1*, and *akt2*) were broadly expressed across cell types, with the highest expression observed in clusters enriched for mTorc1/2 components (Fig. 3*B*), consistent with their role as upstream regulators of mTOR signaling. Expression of insulin/IGF pathway genes revealed cell type specific patterns: as seen during zebrafish fin regeneration (28), *igf2*, an insulin-like growth factor upstream of mTOR signaling (29-32), was selectively upregulated in CT cells, whereas the genes encoding its receptor (*igf1r*), intracellular adaptors (*irs1* and *irs2*) and extracellular modulators (*igfbp2, igfbp3* and *igfbp6*) were most highly expressed in basal epidermal cells and in proliferating CT cells. This complementary expression pattern suggests that IGF signaling may mediate communication between CT cells and the overlying wound epidermis during regeneration. Finally, inflammatory signaling genes *il1b* and *il1r1* were predominantly expressed in myeloid cells, consistent with activation of immune programs that are known to intersect with mTOR regulation. Collectively, these results indicate that mTOR signaling during fin regeneration integrates distinct upstream cues, including IGF-mediated growth factor signaling between connective tissue and epidermal cells, and lysosomal nutrient and inflammatory inputs in myeloid cells.

To gain further insight into the location of mTOR pathway activation in relation to the injury site, we leveraged publicly available spatial transcriptomics datasets corresponding to the same regenerative stages analyzed by snRNA-seq (21) (Fig. 3*C* and *D*). As a readout consistent with mTOR pathway activation, we first examined changes in expression of translation-related genes during fin regeneration, as mTOR signaling regulates transcription of ribosomal RNA genes (33). To this end, we curated a set of 12 genes encoding ribosomal proteins and ribosomal RNA components (*npm1a, ncl, fbl, eif4g1a, eif2s1b, rps6, rps3, rps14, rpl11, rplp0, eef2b, eef1b2*). Uniform Manifold Approximation and Projection (UMAP) analysis revealed that expression of this translation gene set was highly active in subsets of cells across all clusters in the uninjured fin (Fig. 3*E*). At 1 dpa, expression was broadly detectable across epidermal cell clusters, before subsiding at later stages (3 and 7 dpa). Spatial transcriptomic analysis further revealed a distal enrichment of this translation gene set at 3 dpa, indicating that cells with elevated expression of translational machinery were preferentially localized near the injury site. Together, these spatial and transcriptional patterns are consistent with localized activation of mTOR-associated translational programs during early fin regeneration. Next, we evaluated *mtor* gene expression. UMAP showed expression of *mtor* throughout fin regeneration across all cell clusters, yet spatial transcriptomics revealed a distal enrichment at 3 dpa (Fig. 3*F*), resembling the pattern observed for the translation gene set. Expression of *igf2* was largely limited to CT cells, strongly upregulated at 1 dpa, and was distally enriched at 3 dpa like *mtor* and the translation gene set (Fig. 3*G*). In contrast, *igf1r* expression was broadly detected across all cell types, but was most pronounced in the basal epidermis clusters (Fig. 3*H*). Spatial transcriptomics revealed localized expression of *igf1r* in the wound epithelium at 1 dpa, which persisted at 3 and 7 dpa. Finally, *igfbp6b*, a key modulator of Igf2/Igf1r signaling (34), was notably upregulated upon injury and expressed in a subset of basal epidermis cells located in the wound epidermis (Fig. 3*I*), suggesting local fine-tuning of IGF signaling at the injury site. Together, these analyses revealed that mTOR signaling during *Polypterus* fin regeneration is regulated in a cell-type-specific and spatially organized manner, involving IGF-mediated signaling between connective tissue and wound epidermis, lysosomal nutrient sensing and inflammatory signaling in myeloid cells, and localized activation of translational programs near the injury site.

### Rapamycin treatment impacts transcription of ribosomal proteins, glycolysis, and myeloid cell gene expression profiles

Having identified the expression profile of mTOR signaling components during fin regeneration, we next sought to determine the specific impacts of mTOR inhibition. First, we assessed whether rapamycin disrupted the upregulation of genes encoding translational machinery components. We curated a comprehensive list of 276 *Polypterus* genes classified into the following categories: ribosomal RNAs (29 genes), cytosolic ribosomal proteins (79 genes), mitochondrial ribosomal proteins (74 genes), tRNA synthetases (30 genes), translation termination and recycling factors (4 genes), translation elongation factors (15 genes), and translation initiation factors (45 genes) (Dataset S3). Consistent with our prior analysis of a smaller translation gene set, most translation machinery categories were sharply upregulated in DMSO-treated animals at 1 dpa, followed by a gradual decrease at 3 and 7 dpa (Fig. 4*A and B*). In contrast, this upregulation was significantly attenuated in rapamycin-treated animals (Fig. 4*A* and *B*; *SI Appendix*, Fig. S3).

**Fig. 4.**
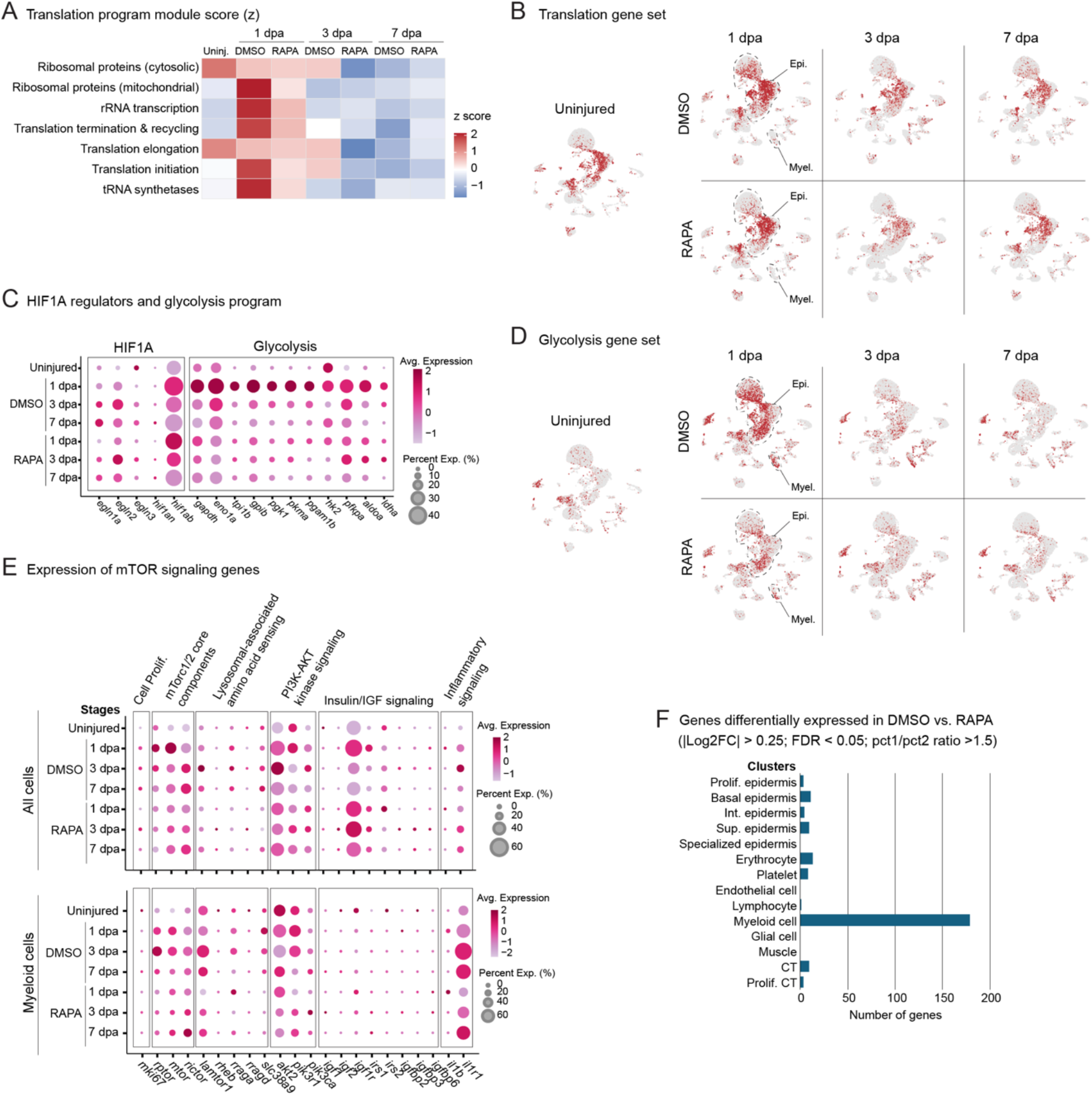
mTOR signaling inhibition impacts genetic programs of translational machinery, glycolysis, and myeloid cells. (A) Heatmap of z-scores for genetic modules of translational machinery gene sets in the uninjured fin and across fin regeneration stages. (B) UMAPs showing enrichment of translation gene set in the uninjured fin and across fin regeneration stages in DMSO- and rapamycin-treated animals. (C) Dot plot showing expression of *hif1ab*, its regulators, and a select set of glycolysis-related genes in the uninjured fin and across fin regeneration stages in DMSO- and rapamycin-treated animals. (D) UMAPs showing enrichment of a glycolysis gene set in the uninjured fin and across fin regeneration stages in DMSO- and rapamycin-treated animals. (E) Dot plot showing expression of mTOR signaling components and modulators in all cells (top) and in myeloid cells (bottom) in the uninjured fin and across fin regeneration stages in DMSO- and rapamycin-treated animals. (F) Bar graph showing differential gene expression between DMSO- and rapamycin-treated fins in consolidated cell clusters from 1, 3 and 7 dpa combined.

We next asked whether mTOR activity is required for activation of hypoxia-associated metabolic programs during fin regeneration, given the well-established role of mTOR signaling in coordinating these processes (1). In both the axolotl limb and *Polypterus* fin, amputation induces HIF1a stabilization and drives a rapid shift toward glycolytic metabolism (21, 35). We therefore examined the temporal regulation of the HIF1a axis, including *hif1a* and its negative regulators (*egln1-3* and *hif1an*), together with a curated set of glycolysis-associated genes, in DMSO- and rapamycin-treated fins across regenerative stages. In DMSO-treated fins, *hif1a* expression increased sharply at 1 dpa, followed by delayed upregulation of its negative regulators at 3 and 7 dpa, consistent with transient HIF1a activation during early regeneration (21) (Fig. 4*C*). Glycolysis genes exhibited a coordinated induction at 1 dpa, followed by progressive attenuation at later stages. Rapamycin treatment did not alter the temporal activation of *hif1a* or its regulatory network. However, we observed little to no upregulation of glycolysis-associated genes at 1 dpa, which instead remained near homeostatic levels throughout regeneration. UMAP visualization of DMSO- and rapamycin-treated datasets revealed that induction of this glycolysis gene set was most prominent in epidermal and myeloid cell clusters (Fig. 4*D*), consistent with mTOR signaling being robustly detected in these cell types during fin regeneration.

To assess the overall impact of mTOR inhibition on its own signaling pathway, we examined expression of genes encoding mTOR pathway components and major upstream regulatory axes. Rapamycin treatment resulted in expression profiles broadly similar to those observed in DMSO-treated animals, both across cell clusters (*SI Appendix*, Fig. S4) and regeneration stages (Fig. 4*E*). Notably, however, we detected a marked attenuation of expression in myeloid cells for genes encoding mTorc1/2 core components and lysosomal amino acid– sensing machinery (*SI Appendix*, Fig. S4). Analysis of myeloid cells across regeneration stages confirmed a pronounced reduction in expression of *mtor, rptor*, and *slc38a9*, the latter encoding a key lysosomal amino acid transporter upstream of mTORC1 activation (36).

Finally, we investigated the global impact of mTOR inhibition on transcriptional programs during regeneration across all cell types by performing differential gene expression analysis. Cells from 1, 3, and 7 dpa were pooled into unified DMSO- and rapamycin-treated conditions, and differential expression was assessed using conservative thresholds (FDR < 0.05; log2 fold change > 0.25). Genes were further filtered to retain those showing substantial differences in expression breadth, requiring a > 1.5-fold difference in the fraction of expressing cells between conditions (pct1/pct2 > 1.5). This analysis revealed that myeloid cells exhibited the strongest sensitivity to mTOR inhibition, with 179 differentially expressed genes (Dataset S4), whereas most other cell types, including epidermal, connective tissue, endothelial, muscle, glial, and erythrocyte populations, showed minimal transcriptional differences (0-13 genes) (Fig. 4*F*). These results indicate that rapamycin does not broadly reprogram cellular transcriptomes during fin regeneration, but instead preferentially affects immune cell populations. Collectively, our findings demonstrate that mTOR inhibition strongly impairs transcriptional upregulation of the translational machinery, disrupts activation of a glycolytic metabolic program, and disproportionately affects myeloid cell gene expression during fin regeneration.

### mTOR inhibition dampens myeloid cell programs associated with regenerative resolution

Regulation of immune cell activation, differentiation, and effector function is a canonical role of mTOR signaling (37), and pharmacological inhibition of mTOR with rapamycin leads to robust immune suppression (38). In myeloid cells, mTOR inhibition has been shown to dampen inflammatory responses while promoting resolution-associated activation states in macrophages (39-42). Given the transcriptional impact we observed in myeloid cells upon rapamycin treatment, we next sought to determine if mTOR inhibition was preferentially attenuating pro-regenerative myeloid programs.

To facilitate pathway annotation, *Polypterus* genes within the set of 179 genes differentially expressed in myeloid cells between DMSO- and rapamycin-treated animals were assigned corresponding human gene symbols. For most genes in this data set, expression levels peaked at 3 dpa in DMSO-treated animals, whereas rapamycin-treated animals showed either homeostatic expression levels or modest induction across regeneration stages (Fig. 5*A*, Dataset S4). Pathway enrichment analysis using the MSigDB Hallmark gene set collection identified significant enrichment for glycolysis and mTORC1 signaling (FDR < 0.05), consistent with our transcriptional analyses showing reduced metabolic and translational activity following mTOR inhibition (Fig. 5*B*). Additional enriched pathways included complement, interferon-alpha and interferon-gamma responses, allograft rejection, and unfolded protein response. Notably, these immune-associated pathways were enriched in the absence of hallmark pro-inflammatory signatures, suggesting that rapamycin-sensitive transcriptional programs correspond to immune-coordinating, interferon-responsive, and metabolically active myeloid states rather than cytokine-dominant inflammatory programs.

**Fig. 5.**
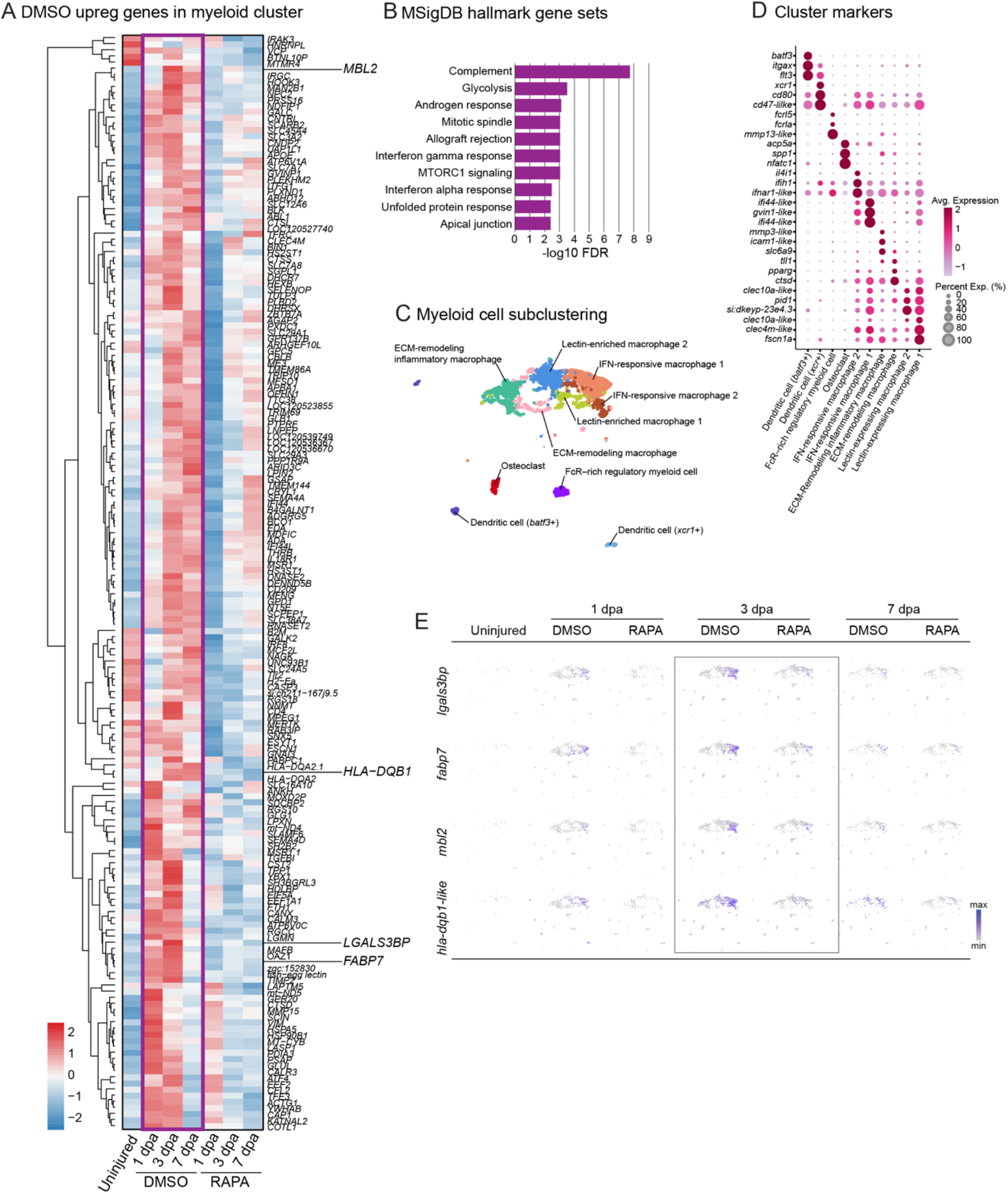
mTOR signaling inhibition disrupts pro-resolution genetic programs in myeloid cells. (A) Heatmap showing human orthologs of Polypterus genes differentially expressed in DMSO-treated relative to rapamycin-treated animals during fin regeneration. (B) Functional classification of signaling pathways enriched in DMSO *versus* rapamycin-treated animals. (C) UMAP showing subclustering of myeloid cells. (D). Dot plot showing expression of myeloid cell cluster markers. (E) UMAPs showing expression of select set of genes upregulated in DMSO-treated *versus* rapamycin-treated animals during fin regeneration.

To gain insight into the myeloid cell states underlying these rapamycin-sensitive pathways, we subclustered the myeloid lineage into nine transcriptionally distinct populations and annotated them based on established marker gene expression (Fig 5*C*). These populations included two dendritic cell (DC) subsets (*batf3*^+^ and *xcr1*^+^ DCs), Fc receptor-rich regulatory myeloid cells, osteoclasts, two interferon-responsive macrophage populations, ECM-remodeling inflammatory macrophages, ECM-remodeling macrophages, and two lectin-expressing macrophage populations. Representative marker genes supporting these annotations are shown in the accompanying dot plot (Fig 5*D*).

Finally, we examined the cellular expression patterns of representative genes from the 179-rapamycin-sensitive gene set. Genes marking lectin-mediated innate recognition (*mbl2*), antigen presentation competence (*hla-dqb1-like*), IFN-associated immune activation (*lgals3bp*), and immunometabolic remodeling (fabp7b) localized preferentially to IFN-responsive and lectin-expressing macrophage states (Fig. 5*E*). Additional genes involved in antigen processing and immune competence (*prss16, ctss, b2m, hla-dqa2-like*) and interferon-responsive activation (*irgc, gvinp1, irf8, ifi44*) exhibit similar localization to IFN-responsive and antigen-presenting myeloid states (*SI Appendix*, Fig. S5). Collectively, these patterns suggest that rapamycin selectively dampens macrophage activation states characterized by innate recognition, immune communication, antigen presentation, and metabolic adaptation, rather than uniformly suppressing myeloid gene expression.

## Discussion

Why salamanders are the only extant tetrapods capable of regenerating complete limbs as adults remains a central unresolved question in regenerative biology. The recent identification of salamander-specific amino acid expansions in the otherwise highly conserved mTOR protein, which render the kinase unusually primed for activation, provided a compelling molecular hypothesis for this exceptional regenerative capacity (17). The absence of such expansions in mammalian mTOR, together with the limited regenerative ability of mammalian limbs (restricted to digit tips) (43), suggested that a hypersensitive mTOR kinase might be required for appendage regeneration. Here, using *Polypterus* fin regeneration as a model system, we show that this is not the case. Our findings demonstrate that a canonical, non-expanded mTOR protein is fully compatible with vertebrate complex appendage regeneration. As observed in axolotl limbs, mTOR activity in *Polypterus* is rapidly induced within hours of fin amputation, with particularly strong activation in the wound epidermis. Moreover, pharmacological inhibition of mTOR signaling completely blocks fin regeneration, establishing mTOR as an essential regulator of regenerative outcome in this species.

SnRNA-seq of DMSO- and rapamycin-treated fins provided a high-resolution view of the cellular consequences of mTOR inhibition. Rather than eliciting a global transcriptional block, rapamycin induced a highly selective response across cell types. Canonical mTOR-dependent processes, including the metabolic shift toward glycolysis and the transcriptional upregulation of genes encoding translation machinery components, were strongly attenuated, consistent with well-established physiological roles of mTOR signaling. Notably, most epidermal, connective tissue, endothelial, muscle, glial, and erythrocyte populations exhibited relatively modest transcriptional changes between treatments. In contrast, myeloid cells displayed a disproportionately large response to mTOR inhibition. Within the myeloid compartment, transcriptional changes were largely confined to antigen-presenting, interferon-responsive, and lectin-associated activation states. These data support a model in which mTOR activity is required to sustain specific myeloid programs associated with regenerative coordination and resolution. This interpretation is consistent with prior work demonstrating that macrophage depletion severely impairs axolotl limb regeneration (44). Thus, the effect of mTOR inhibition in myeloid states during *Polypterus* fin regeneration may explain, at least in part, the severe impact in fin outgrowth observed upon rapamycin treatment.

Collectively, our results support an ancestral role for canonical mTOR signaling in coordinating the translational, metabolic, and immune components of vertebrate appendage regeneration. Multiple lines of evidence indicate that the capacity for limb regeneration was present in early tetrapods and inherited from fish ancestors, including regenerative pathologies observed in fossil amphibians predating modern salamanders (45), the occurrence of complete fin regeneration in members of all major bony fish clades (46), and the deployment of conserved genetic programs during fin and limb regeneration (21, 46, 47). In this context, our findings align with a model in which canonical mTOR signaling represents a core component of the ancestral regenerative toolkit, whereas hypersensitive mTOR activity constitutes a derived salamander-specific adaptation. While enhanced mTOR activity was likely not required for fin or limb regeneration in aquatic ancestors, the transition of early tetrapods to semi-terrestrial environments may have imposed new constraints on regenerative success. For example, rapid wound closure in air-exposed tissues may have become essential to prevent dehydration. The evolution of mechanisms that accelerate wound epidermis formation and closure may therefore have played a key role in preserving regenerative capacity in urodele amphibians while it was lost in other tetrapod lineages.

## Materials and Methods

### Animal work

Juvenile *Polypterus* (*P. senagalus*) were obtained from commercial vendors, maintained in individual tanks in a recirculating freshwater system at 27-28 °C and a day-night cycle as 12 h light/12 h dark, and used in accordance with the approved Louisiana State University (LSU) IACUC protocol IACUCAM-25-047. Before surgical procedure, fish were anesthetized in 0.05% MS-222 (Sigma), diluted in fish system water. Pectoral fins of *Polypterus* fish ranging from 7 to 10 cm were bilaterally amputated across the proximal endoskeleton using a sterile scalpel blade. For uninjured fin samples, a portion of the amputated fins encompassing the endoskeletal elements was sampled. Regenerating fins were harvested from animals treated with DMSO (0.1% in fish system water), rapamycin (2.5 μM) or INK128 (0.5 μM) for subsequent experiments. All animals were euthanized according to our approved IACUC protocol (0.25% MS-222 in the appropriate animal media).

### Alignment of mTOR proteins

The following full-length mTOR proteins were used for sequence alignment and comparisons: *Homo sapiens* (NP_001373429), *Ambystoma mexicanum* (XP_069466034.1), *Polypterus senegalus* (XP_039611786.1), *Danio rerio* (NP_001070679.3), *Pan troglodytes* (XP_016809439.2), *Mus musculus* (NP_064393.2), *Canis lupus* (XP_005618060.1), *Ovis aries* (XP_027831352.1), *Ornithorhynchus anatinus* (XP_028920355.1), *Meleagris gallopavo* (XP_010721303.1), *Gallus gallus* (WWQ02357.1), *Thamnophis elegans* (XP_013920889.1), *Anolis carolinensis* (XP_062821935.1), *Crocodylus porosus*, (XP_019397232.1), *Chrysemys picta* (XP_008165456.1), *Xenopus tropicalis* (XP_031761371.1), *Bufo bufo* (XP_040286249.1), *Microcaecilia unicolor* (XP_030041416.1), *Geotrypetes seraphini* (XP_033778669.1), *Pleurodeles waltl* (XP_069097339.1), *Protopterus annectens* (XP_043919153.1), *Latimeria chalumnae* (XP_064423891.1), *Oryzias latipes* (XP_023812076.1), *Carcharodon carcharias* (XP_041061752.1). Full-length mTOR protein sequences from the indicated species were retrieved from NCBI and aligned using Clustal Omega v1.2.4 (48) with default parameters. The resulting multiple sequence alignment was visualized using MView (49).

### mTOR HEAT domain 3D modeling and structural alignment

Protein structures were predicted using ColabFold (25) with the AlphaFold2 (26) alphafold2_ptm model. mTOR protein sequences from human, zebrafish, *Polypterus*, and axolotl were used as input. For each sequence, five models were generated with six recycles and a fixed random seed. Multiple sequence alignments were obtained using the default ColabFold MMseqs2 server. Models were ranked based on pLDDT scores, and the top-ranked structure was used for downstream analyses. Pairwise structural alignments of the HEAT domain region were performed in UCSF ChimeraX (50) using the Matchmaker tool, which generates a sequence alignment prior to iterative least-squares superimposition. Overall Root Mean Square Deviation (RMSD) values were obtained for all pairwise combinations of the four species.

### Protein expression analysis via western blot

To assess mTOR pathway activity during fin regeneration and its inhibition by rapamycin, protein lysates were prepared from *Polypterus* pectoral fins previously amputated at the endoskeleton level and harvested at the indicated time points (2 hpa, 1 dpa, and 3 dpa) from animals treated with DMSO (0.1%) or rapamycin (2.5 μM). For the uninjured stage, a segment of an intact fin, encompassing the endoskeleton elements, was sampled. For each biological replicate, the dissected pair of fins were quickly minced in a petri dish on ice using a scalpel blade. The minced fins were homogenized in 300 uL of T-PER™ Tissue Protein Extraction Reagent (Thermo Scientific # 78510) containing 1X Halt Protease Inhibitor Cocktail (Thermo Scientific # 87786). Samples were kept in ice all the time and homogenized in Eppendorf tubes using a disposable pellet pestle. After homogenization, samples were centrifuged at 10,000 xg for 5 minutes. Cleared supernatant was transferred to new tubes and stored at -80C. Total protein concentration was determined by BCA assay (Pierce™ BCA Protein Assay Kit, Thermo Scientific #23227). Equal amounts of protein (20 μg per lane) were denatured by incubation at 70 °C in a sample buffer containing 1X of the Bolt LDS sample buffer (Thermo Scientific # B0007), 1X of the Bolt Sample Reducing Agent (Thermo Scientific # B0009), and resolved by electrophoresis in a precast 4-12% Bolt™ Bis-Tris Plus Mini Protein Gel (Thermo Scientific # NW04122BOX or NW04125BOX) in 1X Bolt MES Running Buffer (Thermo Scientific #B0002). Proteins were transferred to PVDF membranes (0.2 mm) using the Mini Blot Module (Thermo Scientific #B1000) wet transfer system, according to the manufacturer’s instructions. Membranes were blocked in 5% BSA in 1X Tris-buffered saline with Tween (10 mM Tris, 150 mM NaCl, 1% Tween) and incubated with primary antibodies diluted in blocking solution at 4°C overnight. Phosphorylation of ribosomal protein S6 was assessed using an anti–phospho-RPS6 antibody (pS6; Ser240/244; Cell Signaling #5364), and total RPS6 was detected using an anti-RPS6 antibody (Cell Signaling #2217). Detection of β-actin by a specific antibody (Cell Signaling # 3700) was used as a loading control. After three washes in 1X Tris-buffered saline with Tween, membranes were incubated with either HRP-conjugated goat anti-mouse (Thermo Scientific #31430) or goat anti-rabbit (Thermo Scientific #31460) secondary antibody for 1 h at room temperature and then washed three times. Membranes were developed using the SuperSignal™ West Pico PLUS Chemiluminescent Substrate (Thermo Scientific #34577) and detection was performed on a ChemiDocMP (BioRad). Band intensities were quantified in ImageJ/Fiji. pS6 and total S6 signals were first normalized to normalized to β-actin to control for loading and then pS6/totalS6 ratio were calculated. For comparisons across conditions and stages, normalized values were expressed relative to the DMSO-treated sample at the corresponding time point (or relative to uninjured control), as specified. The number of biological replicates (n=3) and statistical tests are reported in the legend.

### Histology and skeletal preparation

Uninjured and regenerating tissues were flash-frozen in Tissue-Tek O.C.T. compound (Sakura) using isopentane (2-methylbutane) in a metal canister placed in liquid nitrogen. Fresh tissues were embedded in appropriate molds. Samples were stored at -80°C and later sectioned into 10 μm thick sections in a Leica CM1520 cryostat. Slides were stored at -80°C until subsequent use. Hematoxylin and Eosin staining was performed as previously described (47). H&E slides were visualized and imaged using a Nikon Eclipse N*i* microscope with a Nikon DS-Fi3 camera. Skeletal preparations of *Polypterus* were performed fins using Alcian blue and Alizarin red staining following established protocols for differential visualization of cartilage and mineralized bone (51).

### Nuclei preparation and single-nucleus RNA-sequencing

For analysis of mTOR inhibition during fin regeneration, snRNA-seq was performed on pectoral fins collected from rapamycin-treated *Polypterus* fish. Uninjured fins and regenerating fins at 1, 3, and 7 days dpa were harvested using sterile spring scissors and scalpel blades. For each condition and time point, pectoral fins from two animals were processed independently through nuclei isolation and subsequently pooled to generate one sample. Two biological replicates were prepared per stage, each consisting of nuclei derived from two animals. Excised fin tissue was minced on ice using spring scissors followed by a razor blade and transferred to 0.3× lysis buffer (Active Motif ATAC-seq kit, #105320, diluted 1:3 in lysis dilution buffer containing 10 mM Tris-HCl pH 7.5, 50 mM NaCl, 20 mM MgCl_2_, and 10% BSA). Tissue suspensions were homogenized in a pre-chilled Dounce homogenizer with 12 strokes and passed through a 40 μm cell strainer. The filtrate was immediately diluted with 4 mL of lysis dilution buffer and centrifuged at 400 × g for 5 minutes at 4 °C using a swinging-bucket rotor. Following centrifugation, nuclei pellets were gently washed with 1 mL of PBS containing 0.5% BSA and 40 U/μL RNase inhibitor (Protector RNase Inhibitor, Millipore-Sigma #3335399001), without disturbing the pellet, and then resuspended in 0.5 mL of the same buffer. At this stage, nuclei isolated from the two animals were combined. Nuclei concentration was determined using a hemocytometer, and aliquots corresponding to 1 × 10^6^ nuclei were pelleted by centrifugation at 400 × g for 5 minutes at 4 °C. Nuclei were fixed and cryopreserved using the Evercode Nuclei Fixation Kit v3 (Parse Biosciences; #NF100, #NF200) according to the manufacturer’s instructions. Fixed nuclei were subsequently processed for library preparation using the Evercode WT v3 kit (Parse Biosciences; #WT100A–D, #WT200), targeting recovery of approximately 7,000 nuclei per sample. Sequencing libraries were generated following the manufacturer’s protocol and sequenced by Novogene on an Illumina NovaSeq X Plus system (paired-end, 150 bp), with an average depth of ∼20,000 reads per nucleus.

### Single-nucleus RNA-seq analysis

Sequencing reads from DMSO-treated (52) and rapamycin-treated samples were aligned to the *Polypterus senegalus* reference genome (assembly ASM1683550v1). Raw snRNA-seq FASTQ files were processed using the Trailmaker™ pipeline (Parse Biosciences; v1.4.0, 2024) via the Trailmaker web platform. Initial processing included sample demultiplexing, barcode error correction, read alignment, and transcript quantification. Gene-level expression counts were compiled into cell-by-gene matrices and automatically forwarded to the Trailmaker Insight module for downstream analysis. Within the Insight module, nuclei failing quality-control criteria were excluded using the default automated filtering workflow, which removes background signal, predicted doublets, and low-quality nuclei characterized by elevated mitochondrial transcript content or low transcript detection. Across samples, the median transcript counts after filtering varied between 1,903 and 5,158, reflecting sample-specific thresholds determined by Trailmaker. The upper cutoff for mitochondrial transcript content was set at 0.5%. Doublet detection and removal were performed using the scDblFinder algorithm, with probability thresholds ranging from 0.58 to 0.84. High-quality nuclei were subsequently normalized, subjected to principal component analysis, and integrated across samples using Harmony. Cell clusters were identified using the Leiden community detection algorithm and visualized in two dimensions using Uniform Manifold Approximation and Projection (UMAP). Cluster-enriched marker genes were identified using the Wilcoxon rank-sum test as implemented in the presto package, comparing each cluster against all remaining nuclei. Data visualization, including heatmaps and UMAP-based gene expression plots, was performed using Trailmaker and the Seurat v5 R package (53) in R using the Seurat FeaturePlot function.

### Pathway enrichment using MSigDB

Functional enrichment was performed using the Molecular Signatures Database (MSigDB) Hallmark gene set collection. P values were corrected for multiple hypothesis testing using the Benjamini–Hochberg method, and pathways with FDR < 0.05 were considered significant. Enrichment results were visualized as bar plots showing −log10(FDR).

## Data Availability

All raw and processed data from snRNA-seq (GSE316120) generated here have been deposited in the Gene Expression Omnibus (GEO). Processed *Polypterus* snRNA-seq data have been made available at the Broad Single Cell Portal (https://singlecell.broadinstitute.org/single_cell) (SCP: 3607).

## Acknowledgments

We thank Joe Fontenot and Emily Kane for the picture of *Polypterus* specimen. This work was funded by start-up funds from Louisiana State University (I.S.), and an NSF-Integrative Organismal Systems (IOS) Enabling Discovery through GEnomics (EDGE) grant (2421117, I.S.).

## Author Contributions

J.F.S. and I.S. designed research; J.F.S., G.L. M.T., C.B., G.B., I.S., A.C.D., and W.B.M. performed research. J.F.S., G.L., L.P., R.G., and I.S. analyzed data; J.F.S. and I.S. wrote the paper.

